# Serum neurofilament light chain predicts stroke outcome and is a potential marker for treatment effects of neural stem cell-derived extracellular vesicles in a rat stroke model

**DOI:** 10.64898/2026.01.28.702334

**Authors:** Megan K. Cannon, Alinde R. Fojtik, Charles M. White, Raymond L. Swetenburg, Steven L. Stice, Sean I. Savitz, Emily W. Baker

## Abstract

Acute ischemic stroke (AIS) remains a leading cause of disability worldwide, and effective treatments are urgently needed beyond reperfusion therapy. Translating preclinical success to clinical impact has been hindered by variability in animal models and the lack of translational biomarkers that predict outcomes across species. To overcome these barriers, we developed a robust rat AIS model optimized for consistency and severity, enabling rigorous therapeutic testing. Additionally, we tested a panel of common clinical serum biomarkers to improve translation from rodents to humans. We demonstrated that serum neurofilament light chain (NfL) -a biomarker widely used in clinical stroke studies-strongly correlated with functional outcomes, establishing a translational link that has not been previously reported in rats. Notably, NfL’s predictive capabilities outperformed infarct volume, a key prognostic factor in moderate and severe strokes, as well as traditional serum biomarkers intercellular adhesion molecule-1 (ICAM-1) and S100 calcium binding protein (S100B). Using this platform, we evaluated the therapeutic impact of neural stem cell-derived extracellular vesicles (NSC EVs), a novel biologic therapy poised for clinical trials, on stroke outcome in our rat AIS model. A three-dose regimen of NSC EV over 48 hours produced the best outcomes in stroked animals evidenced by smaller infarct volume, improved neurologic score, and reduced serum NfL, although single-dose and two-dose regimens were both effective at some endpoints. These findings not only validate NfL as a cross-species biomarker but also provide critical dosing insights for NSC EV therapy, accelerating the path from bench to bedside for AIS treatment.

## Introduction

While reperfusion therapy is the gold standard intervention for acute ischemic stroke (AIS), less than half will be functionally independent at Day 90 even when complete cerebral reperfusion is achieved [1-3]. Furthermore, reperfusion therapy does not address the complex cascade of cellular and molecular events triggered by ischemia. Additional treatment options are an urgent, global unmet medical need. Neural stem cell-derived extracellular vesicles (NSC EVs) are a novel biopharmaceutical product with powerful, multi-mechanistic target biology that dampen the inflammatory environment after stroke which inhibits infarct growth and worsening functional impairment [4-6]. Additionally, NSC EV provides unique advantages to cell-based therapies due to its manufacturing scalability, ease of administration, and the ability to cross the blood brain barrier and home to the infarct site [7]. While other biologic therapies including NSC EV show promise in stroke models, it is challenging to translate preclinical results to clinical trials.

Identifying reliable biomarkers that translate from preclinical to clinical studies is a gap that hinders new stroke therapies. Robust and consistent blood-based biomarkers could aid diagnosis, treatment, and prognosis of AIS, while being less expensive and more accessible than neuroimaging [8]. Establishing a biomarker that is mechanistically linked to the investigational drug is critical for clinical trial success, as it enables highly quantitative pharmacological assessment and strengthens the biological rationale for therapeutic efficacy [9]. There have been promising candidates for determining AIS prognosis as well as differentiating between stroke types [10, 11]. Recent clinical studies of anti-inflammatory and neuroprotective investigational drugs for AIS have measured serum levels of pro-inflammatory mediators, such as interleukin-6 (IL-6), interleukin-1 beta (IL-1β), and tumor necrosis factor-alpha (TNFα) [12, 13]. However, no recent trials have successfully met their primary endpoint after Phase 2 despite promising preclinical blood-based biomarker data [12, 13]. This could possibly be due to species differences in post-AIS inflammatory signaling and cell death mechanisms [14]. The dynamic changes in the serum concentration of these proteins in AIS patients, the temporal pattern of which may vary based on stroke severity and/or type, could also contribute to a lack of meaningful readouts.

A promising biomarker for AIS prognosis is neurofilament light chain (NfL), a well-established marker for neuroaxonal injury or degradation [15, 16]. Elevated levels of NfL have been documented in neurological diseases including multiple sclerosis [17], amyotrophic lateral sclerosis (ALS) [18], traumatic brain injury (TBI) [19], and Alzheimer’s disease [20-22]. Clinical studies have shown a strong association between elevated NfL concentration, larger cerebral infarct volume, and poor functional outcomes in AIS patients, demonstrating its potential as a robust serum biomarker and neuroprotective target for AIS [23-29]. Despite these findings, few preclinical studies have evaluated serum NfL levels after stroke; both studies have been in mice and only one measured NfL levels longitudinally [30, 31]. A robust rat transient middle cerebral artery occlusion (tMCAO) model presents an opportunity to develop a biomarker that can directly bridge preclinical data to clinical endpoints.

In this study we addressed these clinical translation roadblocks by first determining that an occlusion time of 150 minutes was optimal for tMCAO, resulting in consistently large infarcts comprising both cortical and subcortical regions, which corresponded with significant functional impairment. Next, we found that serum levels of NfL outperformed S100 calcium binding protein (S100B) and intercellular adhesion molecule-1 (ICAM-1) with respect to prognostic value for AIS severity in our tMCAO model. Lastly, we demonstrated the therapeutic potential of NSC EV at 1-, 2-, and 3-doses with improvements in neurologic deficit, infarct volume, and serum NfL. These results demonstrate that the pharmacological effect of a novel investigational biologic product, NSC EV, can be robustly quantitated by changes in serum NfL levels in a preclinical tMCAO model. Furthermore, these results provide proof-of-concept that serum NfL can be utilized as a biomarker to optimize dose paradigms in the preclinical setting to be translated to the clinic, offering a new strategy to rigorously assess promising investigational drugs and bring new treatment options to stroke patients.

## Methods

### Experimental groups

This study included two experiments: (1) development of the surgical model by comparing the effect of occlusion times on body weight, neurologic deficit score, and cerebral infarct volume, and (2) evaluating the effect of NSC EV on body weight, neurologic deficit score, cerebral infarct volume, and serum levels of NfL, S100B, and ICAM-1. For Experiment 1, rats were placed into the following groups: (1) 90 minute occlusion (*n =16*), (2) 120 minute occlusion (*n=16)* and (3) 150 minute occlusion (*n=17)*. For Experiment 2, rats were placed in the following groups (*n* = *7-20*): (1) untreated, (2) one NSC EV dose, (3) two NSC EV doses, or (4) three NSC EV doses.

### Animals

All experimental protocols performed in this study were reviewed, approved, and complied with the guidelines established by the University of Georgia Institutional Animal Care and Use Committee. All studies utilized adult male Wistar rats (220-270g) to minimize biological variability. Rats were obtained from Charles River Laboratories (Wilmington, MA) and allowed to acclimate to the facility for one week prior to surgery. Rats were pair-housed prior to surgery, and individually housed after surgery, on a 12hr light/dark cycle and given food and water ad libitum.

### Transient middle cerebral artery occlusion (tMCAO)

Transient focal ischemia was induced by occlusion of the right middle cerebral artery (MCA) using the silicon-coated suture insertion method [32, 33]. Briefly, rats were anesthetized with isoflurane inhalation mixed with oxygen. All rats were subcutaneously administered Ropivacaine (3mg/kg) for local analgesia to the incision site and 0.65mg/kg sustained release Buprenorphine (Ethiqa XR®) for systemic analgesia. Throughout the procedure, body temperature was maintained at 37°C using a heating pad. A midline neck incision was made, and the right internal, external, and common carotid arteries (ICA, ECA, CCA) were exposed. After cutting a small hole in the ECA, a silicon coated monofilament suture (Doccol Corporation 403756) was advanced through the ICA lumen to the MCA, until resistance was felt. The surgical incision was closed and animals recovered in their home cage under a heat lamp. Proper monofilament placement, resulting in cerebral ischemia, was confirmed by assessing abnormalities in front paw flexion and circling behavior prior to reperfusion. Animals were excluded if they displayed normal behavior (neurologic score of 8) during these tests. After the MCA was occluded for 90, 120, or 150 minutes for Experiment 1, and 150 minutes for Experiment 2, rats were re-anesthetized, and the filament was withdrawn from the ICA to restore cerebral perfusion. Animals were excluded from the study if they scored normal (neurologic score of 8) approximately 5 hours after stroke (Day 0) or if they reached humane endpoint prior to Day 3. Rats were weighed prior to surgery and daily on Days 0-3. Animals reaching ≥30% body weight loss were euthanized.

### NSC EV formulation and intravenous administration

Cryopreserved vials of the neural stem cell (NSCs) were thawed and expanded in bulk in suspension bioreactors until the target volume of cell culture conditioned media containing EVs was generated. Next, the conditioned media underwent a series of tangential flow filtration (TFF) and chromatography steps to separate and concentrate the EVs. The EVs underwent Nanoparticle Tracking Analysis (NanoSight, Malvern Panalytical), single-particle interferometric reflectance imaging (Unchained Labs) and protein quantification (ThermoFisher) (**Supp. Fig. 1**). The final NSC EV preparation was made by diluting the EVs with formulation buffer to the target concentration. The bulk NSC EV preparation was filled into sterile polypropylene vials as individual doses (2.7e11 particles per kilogram body weight) and stored at -20°C. On the day of dosing, NSC EV vials were thawed at 4°C. Rats were anesthetized with isoflurane and administered NSC EV (460µl) intravenously via the lateral tail vein.

**Supp. Fig. 1.**
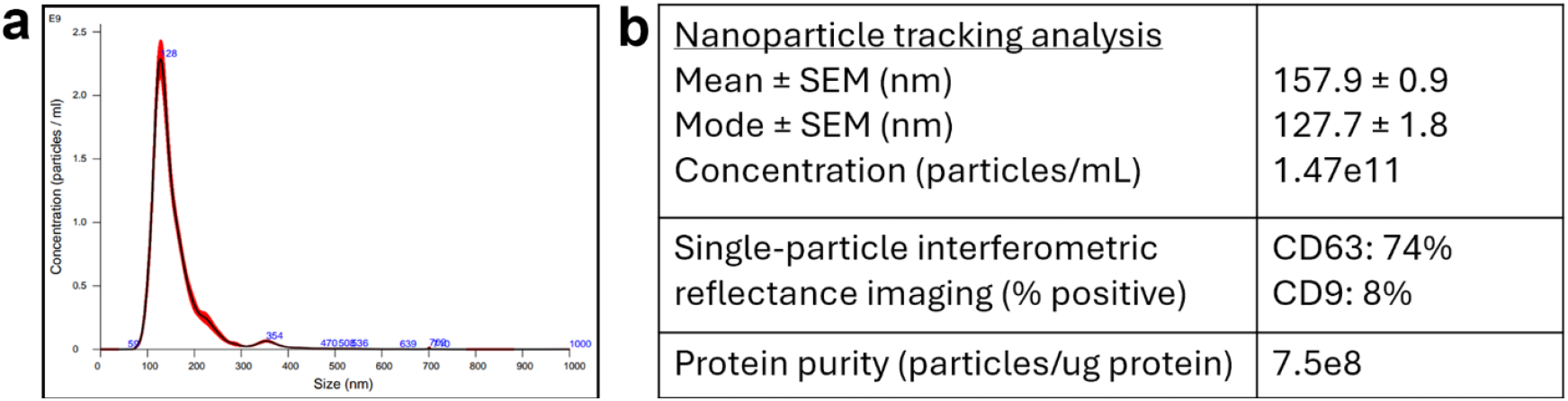
(**a**) NSC EV characterization nanoparticle tracking analysis and (**b**) size, concentration, protein marker and purity specifications.

### Neurologic deficit assessment

All rats were assessed for neurological deficit using a modified version of the Bederson test [34] at 4 hours after stroke reperfusion (i.e., Day 0) as well as Days 1 and 3 post-stroke. Front paw flexion, back paw flexion, circling behavior, and resistance to push were scored on a scale of 0-2, where 0 indicates severe deficit, 1 indicates moderate deficit, and 2 is normal. A total score of 0-8 was then calculated. For receiver operating characteristics (ROC) curves, a good outcome was defined as a neurologic score of 6 or greater on Day 3, while a poor outcome was defined as a neurologic score of 5 or lower. Animals with poor outcomes displayed significant impairment that disrupted quality of life such as trouble accessing and consuming standard rodent chow. Animals with a good outcome exhibited minimal impairment of everyday functioning.

### Serum processing and measurement of serum biomarkers

Blood samples were collected from the ventral tail artery of all animals on Days 1, 2, and 3 post-stroke. Samples were placed into serum separator tubes, kept at room temperature for 30 minutes to allow for coagulation then centrifuged at 4000 x g for 10 minutes to separate serum. NfL (UMAN Diagnostics), ICAM-1 (Invitrogen) and S100B (Invitrogen) proteins were quantified according to the manufacturer’s instructions. All plates were analyzed using the SpectraMax id5 Multimode Microplate Reader (Molecular Devices).

### Evaluation of infarct volume

All animals were euthanized on Day 3 by transcardial perfusion with phosphate buffer saline. The whole brain was extracted immediately then sliced coronally into 6 consecutive 2mm sections using a rat brain slicer matrix (Zivic Instruments). The extent of cerebral infarct was evaluated by placing sections in 2% 2,3,5-Triphenyltetrazolium chloride (TTC, Sigma-Aldrich) and incubating for 10 minutes at 37°C. The area of infarct (i.e., colorless tissue), ipsilateral hemisphere, and contralateral hemisphere was measured for each section using Fiji ImageJ 2.14 (NIH, Bethesda, USA). The areas for each section were multiplied by slice thickness (2mm) then summed into a total volume. Total hemispheric lesion volume with edema correction (%HLVe) was calculated as a percentage using the following formula[35]:

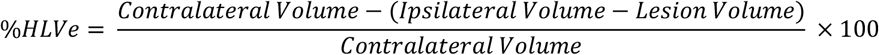

### Statistical Analyses

Line graphs were illustrated with mean ± standard error of the mean (SEM). Violin plots were illustrated with median and quartiles. All data were analyzed using GraphPad Prism 10.4.2 and/or JMP 18.0.2. Statistical analysis of neurologic deficit and body weight was performed using two-way analysis of variance (ANOVA) and post-hoc Tukey’s Pair-Wise comparisons. Pearson correlation and ROC curves were generated and analyzed with JMP. For ROC curves, a good outcome was defined as a neurological deficit score of 6 or greater.

## Results

### Increasing occlusion time to 150-minutes results in higher stroke-induced weight loss and infarct volume, as well as decreased incidence of subcortical infarcts

**Supp. Fig. 2.**
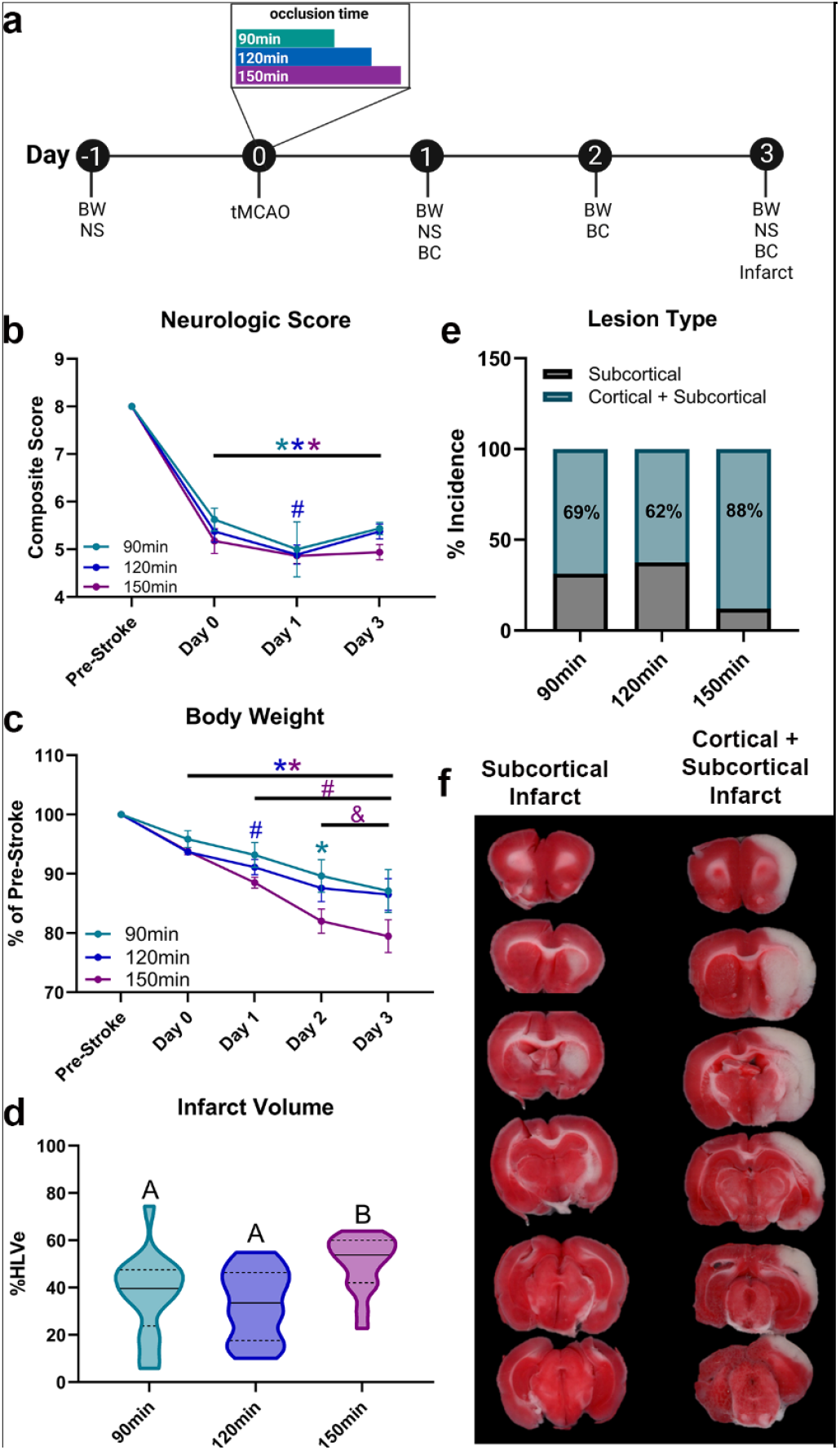
Increasing occlusion time to 150-minutes results in higher stroke-induced weight loss and infarct volume, as well as decreased incidence of subcortical infarcts. (**a**) Experimental outline showing pharmacological time points relative to tMCAO surgery. (**b**) Neurologic score collected pre-stroke and Day 0, 1, and 3. (**c**) Body weight data collected pre-stroke and Day 0, 1, 2, and 3. (**d**) Quantification of cerebral infarct volume on Day 3. (**e**) Occurrence of subcortical vs. cortical + subcortical lesion types for all occlusion times. (**f**) Representative images of TTC-stained coronal brain slices of subcortical only infarct (**left**) and infarct that extends to cortical and subcortical regions (**right**). Data for **b, c**, and **d** expressed as Mean ± SEM; Data for **e** expressed as a percentage of cortical + subcortical strokes. * indicates *p* < 0.05 compared to pre-stroke; # indicates *p* < 0.05 compared to Day 0; & indicates *p* < 0.05 compared to Day 1. For panel **a, *BW:*** body weight, ***NS:*** neurologic score, and ***BC:*** blood collection. For panel **d**, levels not connected by the same letter are significantly (*p* < 0.05) different.

It is critical for laboratories to optimize their model and endpoints prior to treatment studies so that it most closely mimics the intended clinical population. We have previously shown that NSC EV effects are positively correlated with stroke severity, with the most robust treatment response observed in animals with poorer prognosis [4]. Therefore, our goal was to rigorously optimize a preclinical tMCAO model with a highly reproducible and severe neural injury. The model optimization study design is summarized in **Supp. Fig. 2a**. Neurologic deficit was measured before stroke and on Day 0, 1, and 3 post-stroke using a modified version of the Bederson test[34]. Neurologic deficit was represented as a composite score between 0-8, where 8 signified no deficit. Neurologic scores for the 90-, 120-, and 150-minute groups were significantly (*p* < 0.05) lower than pre-stroke on Day 0, Day 1, and Day 3 (**Supp. Fig. 2b, Table 1**). The 120-minute group had significantly (*p* < 0.05) lower neurologic scores on Day 1 compared to Day 0 (**Supp. Fig. 2b, Table 1**). These results confirm stroke induction for all occlusion times which is evident by significantly decreased neurologic scores persisting through Day 3.

**Table 1.**
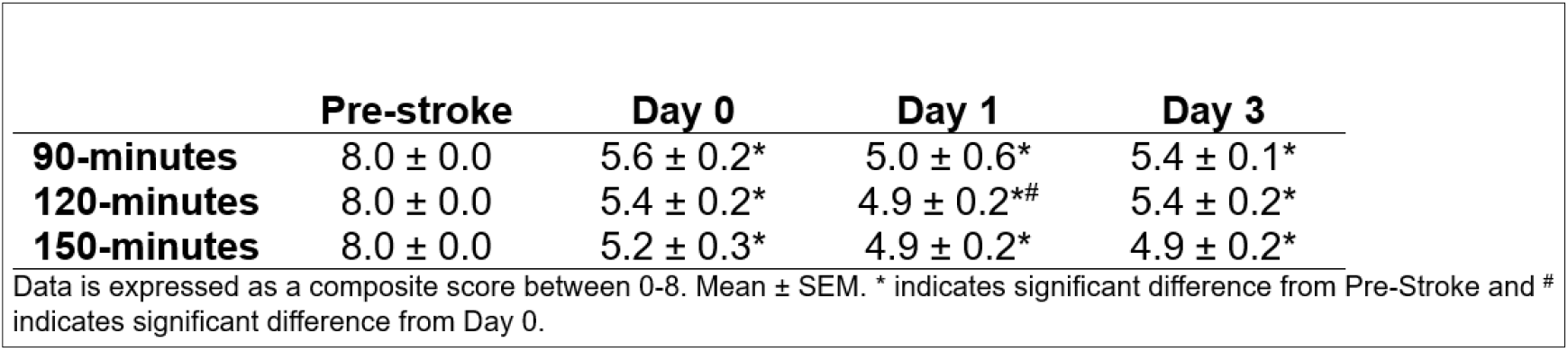
Neurologic deficit score between occlusion times.

Body weight, represented as a percentage of pre-stroke, was measured for 90-, 120-, and 150-minute groups before stroke and on Day 0, 1, 2, and 3 post-stroke. Body weights in the 120- and 150-minute groups were significantly (*p* < 0.05) lower than pre-stroke on Day 0, Day 1, Day 2, and Day 3 (**Supp. Fig. 2c, Table 2**). In the 90-minute group, there was no difference in body weight between pre-stroke and Day 0, Day 1, or Day 3, but body weight was significantly (*p* < 0.05) lower on Day 2 compared to pre-stroke (**Supp. Fig. 2c, Table 2**). In the 120-minute group, body weight on Day 1 was significantly (*p* < 0.05) lower compared to Day 0 (**Supp. Fig. 2c, Table 2**). For the 150-minute group, body weights on Day 1, Day 2, and Day 3 were significantly (*p* < 0.05) lower than Day 0 (**Supp. Fig. 2c, Table 2**). Furthermore, in the 150-minute group, body weights on Day 2 and Day 3 were significantly (*p* < 0.05) lower than on Day 1 (**Supp. Fig. 2c, Table 2**). Taken together, these results show that occlusion time is associated with the degree of stroke-induced weight loss, with the 150-minute group demonstrating greater weight loss over Days 0-3 compared to the 90-minute and 120-minute groups.

**Table 2.** Body weight between occlusion times.

|  | Pre-stroke | Day 0 | Day 1 | Day 2 | Day 3 |
| --- | --- | --- | --- | --- | --- |
| **90-minutes** | 100.0 ± 0.0 | 95.9 ± 1.4 | 93.2 ± 2.1 | 89.6 ± 2.7* | 87.1 ± 3.6 |
| **120-minutes** | 100.0 ± 0.0 | 93.7 ± 0.5* | 91.1 ± 1.3*# | 87.6 ± 2.3* | 86.5 ± 2.7* |
| **150-minutes** | 100.0 ± 0.0 | 93.8 ± 0.6* | 88.5 ± 0.9*# | 82.0 ± 2.0*#& | 79.5 ± 2.8*#& |

Data is expressed as a percentage of Pre-stroke body weight. Mean ± SEM. \* indicates significant difference from Pre-Stroke, # indicates significant difference from Day 0 and & indicates significant difference from Day 1.

Cerebral infarct volume, represented as a percentage of edema corrected hemispheric lesion volume (%HLVe), was measured on Day 3 through TTC staining. Animals in the 150-minute group had significantly (*p* < 0.05) higher infarct volume than both 90- and 120-minute groups (**Supp. Fig. 2d, Table 3**). Furthermore, we categorized the data by infarct type, i.e., whether the cerebral infarct spanned both cortical and subcortical regions or subcortical regions only, and found that the 150-minute group had higher incidence of infarcts spanning both the cortical and Overall, these data demonstrate that a 150-minute occlusion time leads to greater infarct volume and reduced incidence of subcortical infarcts compared to the 90- and 120-minute occlusion times. Since NSC EV has been shown to confer greater benefit in severe strokes [36], these results identify an occlusion time of 150 minutes as the optimal model for subsequent treatment studies.

**Table 3.** Infarct volume between occlusion times.

|  | **Day 3** |
| --- | --- |
| **90-minutes** | $37.3 \pm 4.3^a$ |
| **120-minutes** | $33.0 \pm 3.7^a$ |
| **150-minutes** | $50.4 \pm 2.8^b$ |

Data is expressed as total hemispheric lesion volume with edema correction (%HLVe). Mean ± SEM. Different letters indicate significant differences between groups.

### Elevated concentration of serum NfL is associated with increased stroke deficit and shows great prognostic value for good neurologic outcome

Univariate correlation analyses were performed to examine the relationship between three serum biomarker concentrations (S100B, ICAM-1, and NfL) with body weight and infarct volume. There was no significant correlation between body weight and S100B or ICAM-1 serum concentration on Day 3 (r = 0.45, p = 0.07 and r = 0.32, p = 0.24, respectively, **Fig. 1a, c**). There was also no significant correlation between infarct volume and serum concentration of S100B and ICAM-1 on Day 3 (r = -0.003, p = 0.99 and r = -0.20, p = 0.47, respectively, **Fig. 1b, d**). These results suggest that serum S100B and ICAM-1 are not robust biomarkers for this AIS model, therefore they were not investigated in subsequent studies.

**Fig. 1.**
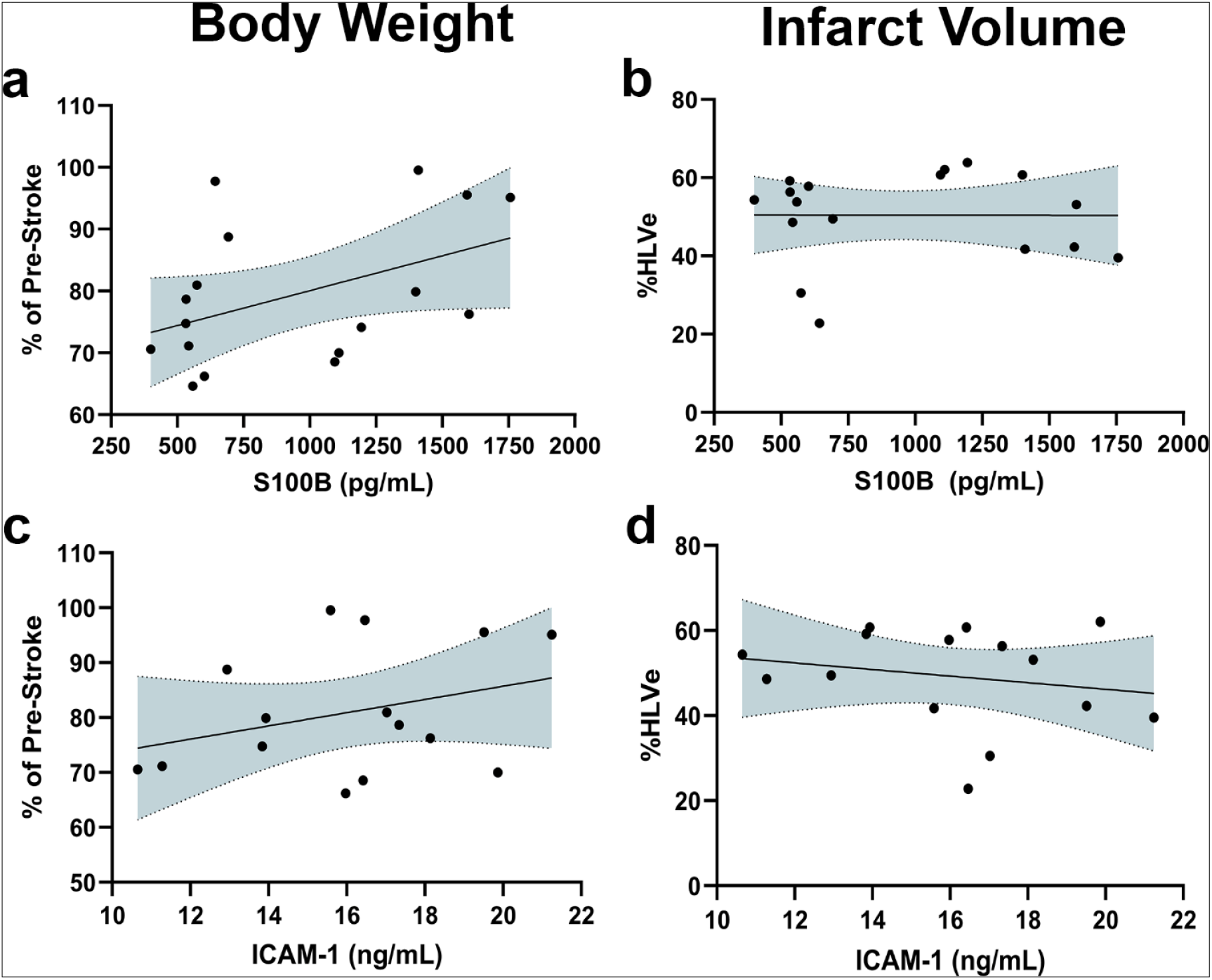
Neither serum S100B nor ICAM-1 are strongly correlated with body weight and infarct volume. Correlation of Day 3 S100B with (**a**) body weight and (**b**) infarct volume. Correlation of Day 3 ICAM-1 with (**c**) body weight and (**d**) infarct volume.

There was a significant correlation between serum NfL concentration and body weight (r = - 0.36, p < 0.05) as well as infarct volume (r = 0.31, p < 0.05) on Day 1 (**Fig. 2a-b, e**). The strength of the correlation between NfL with body weight (r = -0.65, p < 0.01) and infarct volume (r = 0.76, p < 0.01) increased considerably on Day 2 (**Fig. 2e; Supp. Fig. 3**). The correlation between NfL and body weight (r = -0.79, p < 0.01) and infarct volume (r = 0.76, p < 0.01) was strongest on Day 3 (**Fig. 2c-e**). To investigate whether serum NfL level at Day 1, 2 or 3, or infarct volume on Day 3 could discriminate between good and poor functional outcomes, we performed ROC analysis. A good outcome was defined as a neurologic score of 6 or greater on Day 3, while a bad outcome was defined as a neurologic score of 5 or lower. Neither Day 1 nor Day 2 NfL ROC analyses could delineate functional outcomes (Day 1: p = 0.164, AUC = 0.628 and Day 2: p = 0.086, AUC = 0.672, **Fig. 2f**). However, Day 3 NfL successfully differentiated between good and poor functional outcomes (p = 0.002, AUC = 0.779, **Fig. 2f**). Notably, Day 3 NfL level was a better predictor of a good outcome than infarct volume (p = 0.0051, AUC = 0.726, **Fig. 2f)**. Infarct volume had a slightly higher sensitivity than Day 3 NfL at its respective cutoff point (84.2% vs 73.7%) but had a much lower specificity (84.0% vs 56.0%). The results show that the relationship between serum NfL concentration and worsening body weight and infarct volume strengthens over time. Further, Day 3 NfL, but not Day 1 or Day 2 NfL, can predict a good outcome to a greater degree than infarct volume in our rat tMCAO model. Taken together, these data suggest that serum NfL is a robust biomarker for AIS severity and can serve as an indicator of functional outcome in both preclinical and clinical settings.

**Fig. 2.**
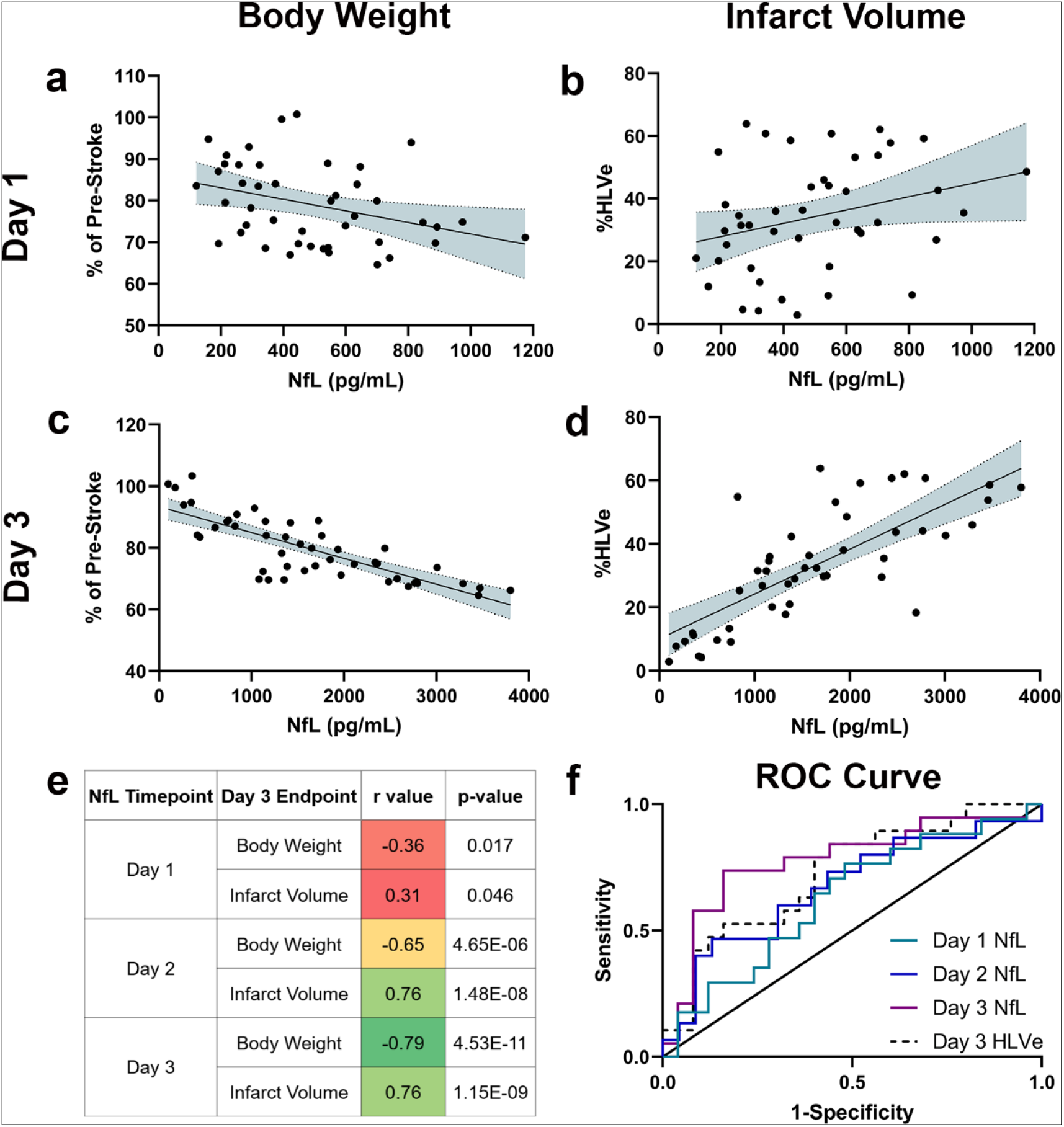
NfL correlates with body weight loss and infarct volume and is predictive of good functional outcomes. Correlation of NfL with (**a**) body weight or (**b**) infarct volume at Day 1. Correlation of NfL with (**c**) body weight or (**d**) infarct volume at Day 3. (**e**) Table showing r value and p-value between NfL and body weight or infarct volume across the three time points. (**f**) Receiver operating characteristic curve for good functional outcomes of Day 1 NfL (green line), Day 2 NfL (blue line), Day 3 NfL (purple line) and lesion size (dashed line).

**Supp. Fig. 3.**
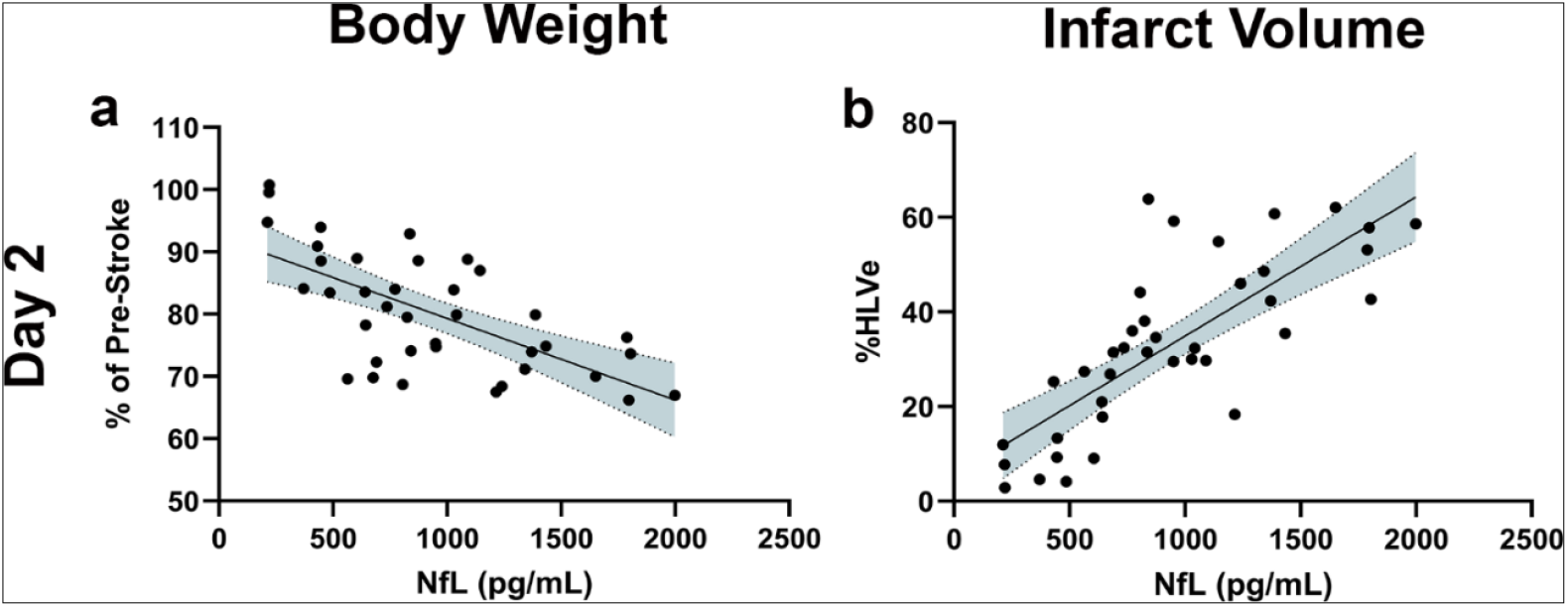
Day 2 serum NfL is correlated with body weight and infarct volume. Correlation of Day 2 NfL with (**a**) body weight or (**b**) infarct volume.

### NSC EV treatment significantly improves stroke-induced weight loss, neurologic deficit, infarct volume, and elevated NfL concentration

We utilized the 150-minute tMCAO model to directly compare the therapeutic effect of three NSC EV dose regimens on stroke outcomes through body weight loss, neurologic score, infarct volume, and serum NfL levels. The dose regimens are outlined in **Fig. 3a**.

**Fig. 3.**
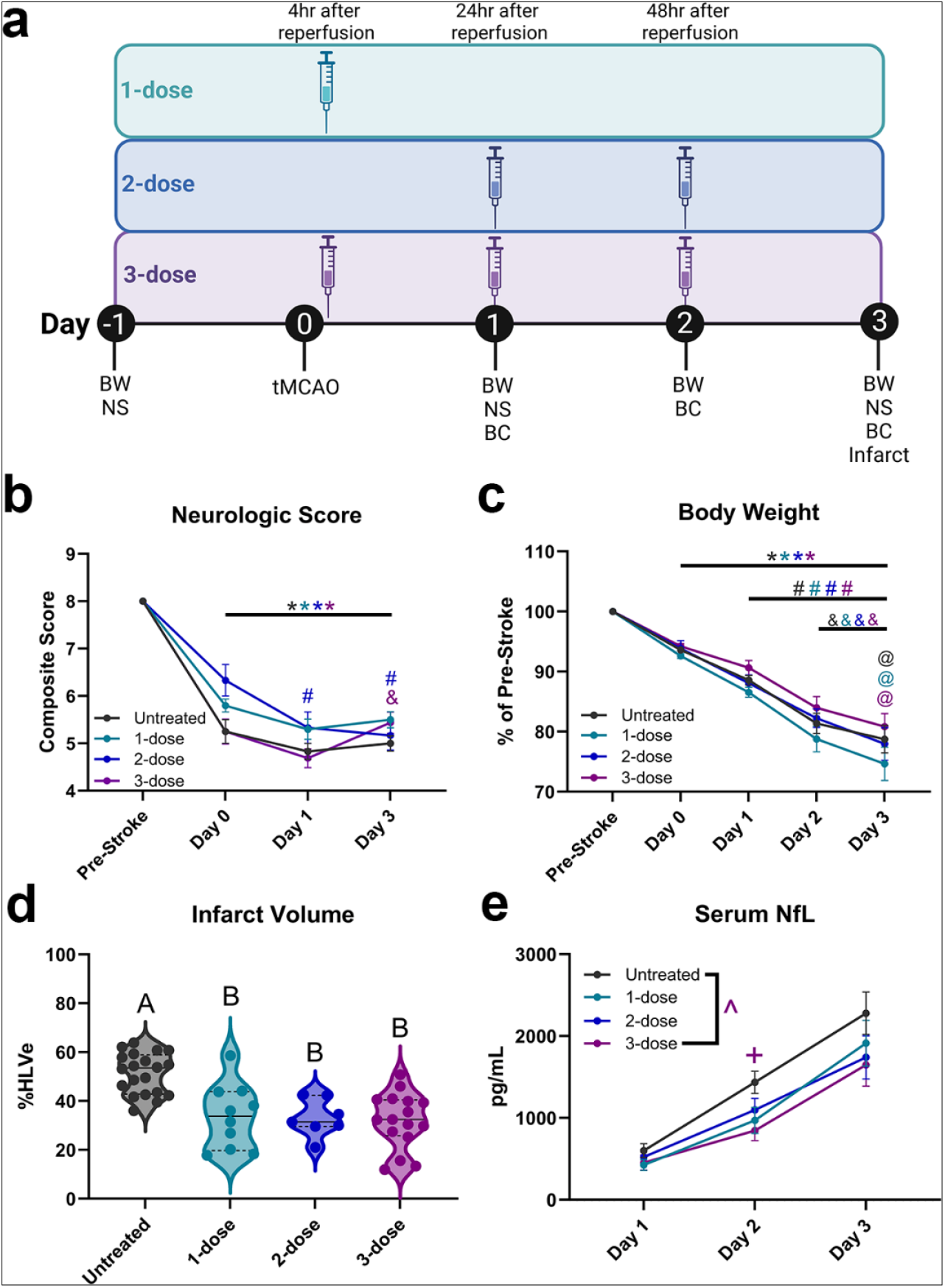
NSC EV treatment significantly improves stroke-induced weight loss, neurologic deficit, infarct volume, and elevated NfL concentration. (**a**) Experimental outline showing treatment schedules and pharmacological timepoints relative to tMCAO surgery. (**b**) Neurologic score collected pre-stroke and Day 0, 1, and 3. (**c**) Body weight data collected pre-stroke and Day 0, 1, 2, and 3. (**d**) Quantification of cerebral infarct volume on Day 3 for all treatment groups. (**e**) Quantification of NfL concentration for all groups on Day 1, 2, and 3. Data expressed as Mean ± SEM. * indicates *p* < 0.05 compared to pre-stroke; # indicates *p* < 0.05 compared to Day 0; & indicates *p* < 0.05 compared to Day 1; @ indicates *p* < 0.05 compared to Day 2. For panel **a, *BW:*** body weight, ***NS:*** neurologic score, and ***BC:*** blood collection. For panel **c**, levels not connected by the same letter are significantly (*p* < 0.05) different. For panel **e**, ^ indicates significant (*p* < 0.05) main effect between untreated vs. 3-

Neurologic scores for the untreated, 1-dose, 2-dose, and 3-dose groups on Day 0, Day 1, and Day 3 were significantly (*p* < 0.05) lower compared to pre-stroke, which confirms stroke induction (**Fig. 3a, Table 4**). Neurologic scores for the 2-dose group on Day 1 and Day 3 were significantly (*p* < 0.05) lower compared to Day 0 (**Fig. 3a, Table 4**). The 3-dose group demonstrated a significantly (*p* < 0.05) higher neurologic score on Day 3 compared to Day 1 (**Fig. 3a, Table 4**). This data suggests that the 3-dose NSC EV regimen led to neurologic improvement between Days 1 and 3.

**Table 4.**
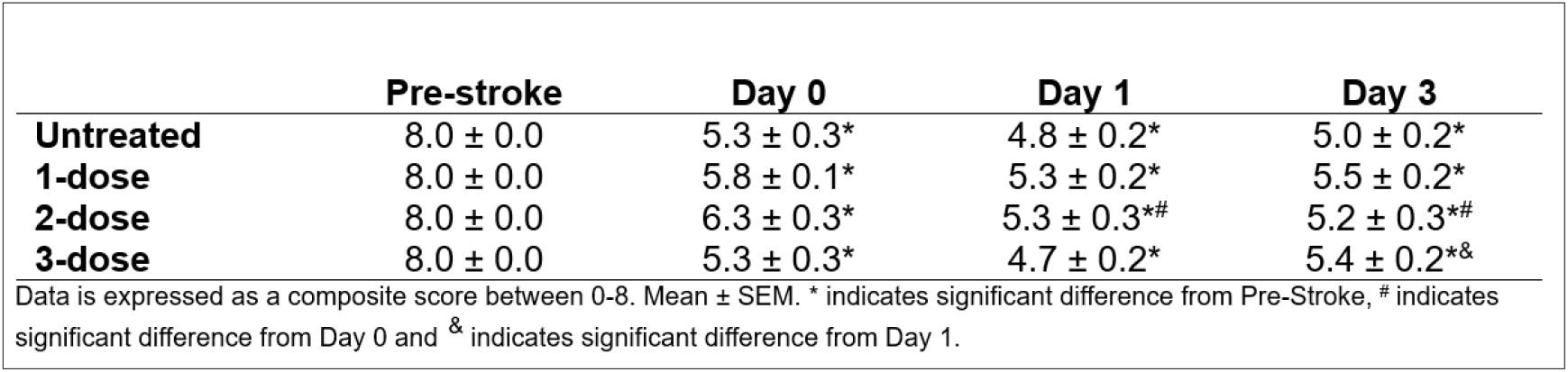
Neurologic deficit score between treatment groups.

Body weights for the untreated, 1-dose, 2-dose, and 3-dose groups were significantly (*p* < 0.05) lower on Day 0, Day 1, Day 2, and Day 3 compared to pre-stroke (**Fig. 3b, Table 5**). In all four groups, body weight was significantly (*p* < 0.05) lower on Day 1, Day 2, and Day 3 compared to Day 0 (**Fig. 3b, Table 5**). In all four groups, body weight was significantly (p < 0.05) reduced on Day 2 and Day 3 compared to Day 1 (**Fig. 3b, Table 5**). The untreated, 1-dose, and 3-dose groups had significantly (*p* < 0.05) lower body weight on Day 3 compared to Day 2 (**Fig. 3b, Table 5**). Together, these data demonstrate that none of the three NSC EV dose regimens mitigated stroke-induced weight loss during the first three days post-stroke.

**Table 5.**
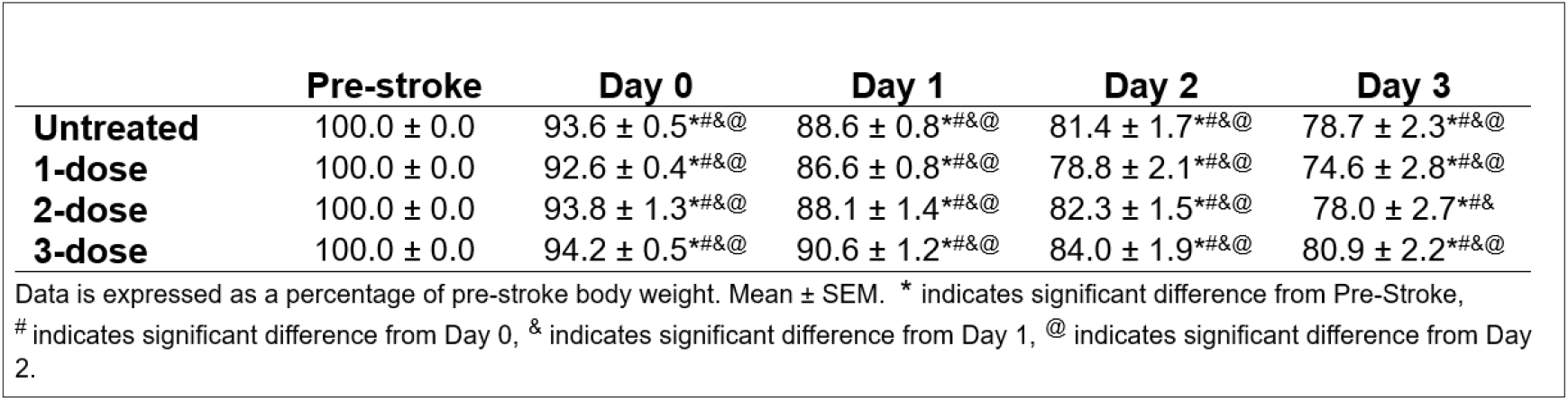
Body weight between treatment groups.

Cerebral infarct volume was measured on Day 3. Infarct volume was significantly (*p* < 0.05) lower in the 1-dose, 2-dose, and 3-dose groups compared to untreated animals (**Fig. 3c, Table 6; Supp. Fig. 4**). These results indicate that volume by Day 3.

**Table 6.** Infarct volume between treatment groups.

|  | Day 3 |
| --- | --- |
| **Untreated** | 51.4 ± 1.9<sup>a</sup> |
| **1-dose** | 33.6 ± 4.2<sup>b</sup> |
| **2-dose** | 33.1 ± 2.9<sup>b</sup> |
| **3-dose** | 32.0 ± 2.8<sup>b</sup> |

Data is expressed as total hemispheric lesion volume with edema correction (%HLVe). Mean ± SEM. Different letters indicate significant differences between groups.

**Supp. Fig. 4.**
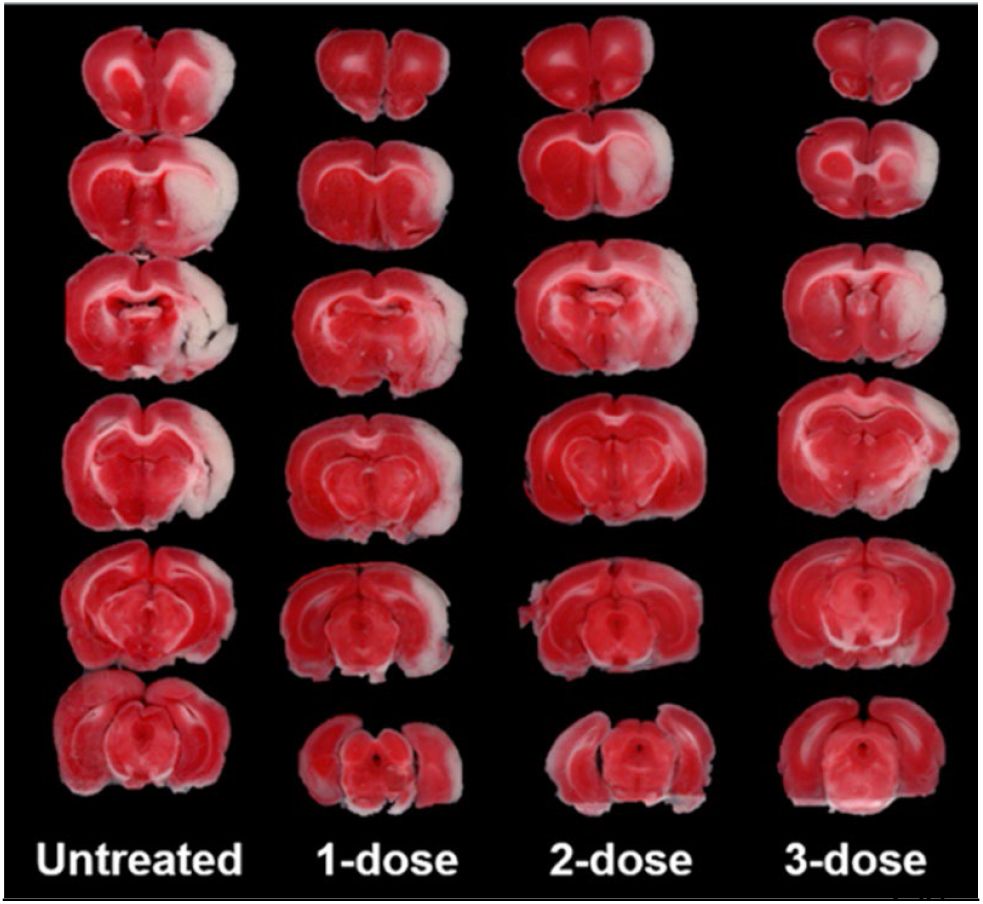
Representative images of coronally sectioned, TTC-stained brain slices for an untreated animals (**Fig. 3c, Table 6;** untreated animal and animals treated with 1-, 2-, or 3-doses of NSC EVs.

Serum NfL concentration (pg/mL) was measured on Days 1, 2, and 3. The 3-dose group had significantly (p < 0.05) lower levels of serum NfL compared to the untreated group on Day 2 (**Fig. 3d, Table 7**). There was also a significant main treatment effect (p < 0.05), where the 3-dose animals had significantly lower serum NfL than the untreated group across all time points (**Fig. 3d, Table 7)**, suggesting that only the full, 3-dose regimen could reduce NfL levels over time.

**Table 7.** Serum NfL concentration between treatment groups.

|  | Day 1 | Day 2 | Day 3 |
| --- | --- | --- | --- |
| **Untreated** | 602.3 ± 84.8 | 1435.5 ± 135.8 | 2279.3 ± 258.6 |
| **1-dose** | 425.2 ± 65.2 | 970.7 ± 159.0 | 1913.8 ± 279.0 |
| **2-dose** | 523.3 ± 88.9 | 1099.3 ± 140.5 | 1740.3 ± 263.7 |
| **3-dose<sup>^</sup>** | 454.6 ± 90.7 | 846.5 ± 121.6<sup>+</sup> | 1649.1 ± 260.1 |

Data is expressed as pg/mL. Mean ± SEM. <sup>^</sup> indicates a main effect significant difference from Untreated. <sup>+</sup> indicates significant difference from Untreated at Day 2.

## Discussion

Novel treatments to aid in stroke recovery after reperfusion are an urgent medical need, however there are many barriers that impede the translation of strong preclinical data to positive clinical trial outcomes. A main objective of our study was to identify a reliable serum biomarker in our rat tMCAO that was responsive to NSC EV treatment effects and could be carried forward to future clinical studies. Two recent clinical studies of anti-inflammatory and neuroprotective investigational products for AIS, ApTOLL and Multistem, measured serum levels of pro-inflammatory mediators as potential biomarkers at both the preclinical and clinical stages. ApTOLL was shown to significantly reduce plasma levels of IL-12p70, interferon-gamma (IFNγ), and IL-6 at 24 hours after permanent MCAO in mice [37]. However, these results did not translate to significant biomarker changes in AIS patients during their Phase 1b/2a clinical study (APRIL) [12]. There are a couple of possible explanations for this lack of translation. First, it is well-documented that pro-inflammatory cytokines are only transiently elevated in serum after stroke, and preclinical rodent models may demonstrate a more rapid temporal pattern compared to human patients [14]. Therefore, it may be difficult to measure a treatment effect at the cytokine’s peak when only a few time points are collected in the sub-acute phase following stroke. Second, while the mice used in the preclinical study underwent permanent occlusion, participants in the APRIL trial underwent endovascular thrombectomy (EVT) and thus were reperfused. This key difference in reperfusion status affects both the magnitude and the timing of the inflammatory cytokine response after stroke [14]. Studies in a rat tMCAO model demonstrated that Multistem significantly reduced serum levels of IL-6 and IL-1β four days after intravenous infusion [38]. These findings successfully translated to the MASTERS trial, in which investigators showed that IL-6, IL-1β, and TNFα were significantly reduced on Day 7, unveiling some of the first data showing that an anti-inflammatory product modulates immune responses after AIS in human patients [13]. A potential advantage to the Multistem program, compared to ApTOLL, is that both the rat model and AIS patients were transiently occluded, better harmonizing stroke pathophysiology between the preclinical model and the clinical population. Overall, the lack of progress in developing a blood-based inflammatory cytokine biomarker has likely discouraged its evaluation in other recent AIS trials including Nelonemdaz and ARG-007 [39, 40].

We evaluated S100B, ICAM-1, and NfL as potential serum-based biomarkers to quantitate NSC EV treatment effects in a rat tMCAO model. A previous study in EVT patients showed that serum concentration of S100B correlated with infarct size, and median S100B concentrations were lower on Day 2 in patients with a favorable outcome compared to an unfavorable outcome [41]. Similarly, previous preclinical studies have shown that serum S100B is positively correlated with brain edema, hemorrhage, infarct volume, and neurological outcome in rat tMCAO models [42, 43]. However, S100B has been shown in the clinic to peak in the first 24 hours followed by a rapid decrease [44], and it loses much of its prognostic value by Day 3 [45]. Likewise, serum levels of ICAM-1, an adhesion molecule known to be upregulated in cerebral blood vessels after stroke to facilitate immune cell infiltration, have been associated with early neurological deterioration and worse outcome in AIS patients [46, 47]. Despite these data, we found that serum levels of S100B and ICAM-1 were poorly correlated with body weight loss and infarct volume on Day 3 post-stroke in the tMCAO rat model. This could be due to the timing of S100B measurement, since S100B peaks in rats at 48 hours after tMCAO while ICAM-1 peaks at 12 hours after stroke onset [42, 48]. Given the 3-dose regimen of NSC EV administered once daily outperformed the 1- and 2-dose regimens, a biomarker with a more durable and robust stroke-induced response is desirable. NfL remains in circulation longer than pro-inflammatory cytokines, S100B, and ICAM-1, making it a promising candidate to monitor AIS prognosis and potential treatment effects more reliably and over a longer period of time [30, 49].

Here, serum NfL was correlated with both infarct volume and body weight loss, with the correlation becoming stronger over Days 1 through 3. NfL, a neuronal cytoskeleton protein, is a blood-based biomarker for severity at hospital admission and functional prognosis in AIS patients [50, 51]. Plasma levels of NfL are significantly elevated in AIS patients compared to healthy controls up to 6 months post-stroke, with Day 7 levels being the best predictor of moderate-to-high stroke (NIHSS ≥5)[49]. However, despite the evidence indicating its utility in clinical AIS, serum NfL in preclinical stroke models remains largely understudied. Plasma NfL was elevated at 24 hours through 7 weeks post-stroke in a mouse permanent distal MCAO with hypoxia model, while plasma NfL was elevated on Day 30 in a mouse tMCAO model [30, 31] However, no prior longitudinal NfL studies have been conducted in a rat stroke model, a species that is widely utilized for AIS drug development. Our longitudinal results align with a recent report indicating that plasma NfL was positively correlated with infarct volume and independently predicted Day 90 Modified Rankin Score (mRS) more effectively than infarct volume in AIS patients [52]. These results corroborate our findings and suggest that NfL could be a robust biomarker for monitoring AIS prognosis and treatment effects and can be used as a benchmark for success in AIS drug development at the preclinical stage.

The 3-dose regimen of NSC EV significantly reduced serum NfL in the rat tMCAO model, indicative of reduced neuronal damage after stroke. NSC EV suppresses inflammatory signaling cascades in the acute post-stroke period, reducing the extent of neural injury, through the activity of therapeutic cargo (miRNAs and proteins) and surface molecules [5, 6]. Cell-based assays and animal stroke models have elucidated several potential direct and indirect pathways by which NSC EV functions to preserve brain tissue and improve stroke outcomes [4-6]. NSC EV possesses AMPase enzymatic function, which is involved in converting adenosine triphosphate (ATP) to anti-inflammatory adenosine, through the activity of CD73 on the EV membrane [53]. Furthermore, NSC EV reduces protein levels of proinflammatory C-C Motif Chemokine Ligand 2 (CCL2) in activated human microglia cells[54]. Reduced CCL2 signaling could lead to a more intact blood-brain barrier and less immune cell infiltration into the CNS, as well as reduced polarization of these immune cells to a pro-inflammatory phenotype [55, 56]. NSC EV mitigates the systemic immune response evident by significant reductions in pro-inflammatory circulating Th17 cells as well as significant increases in anti-inflammatory T regulatory cells and M2 macrophages in a rodent thromboembolic stroke model, leading to reduced infarct volume and preserved sensorimotor function [6]. Together, these mechanisms are thought to reduce infarct growth evident by reduction in circulating NfL.

We found that all three NSC EV dosing regimens significantly reduced infarct volume at Day 3 post-stroke. This effect is notable given the timing of treatment initiation: whereas the 1-dose and 3-dose regimens began 4 hours after reperfusion, the first dose of the 2-dose regimen was delayed until 24 hours post-reperfusion. Although reduced infarct volume in the 2-dose group did not correlate with improvements in neurologic score or body weight, future studies will focus on optimizing functional assessments to confirm NSC EV–mediated effects when treatment is initiated at later time points. The ability of NSC EV to reduce infarct volume despite delaying the first administration to 24 hours suggests potential utility in a broader patient population, including individuals who have missed the therapeutic window for tissue plasminogen activator [57].

In conclusion, we show for the first time in a rat stroke model that serum NfL is highly correlated to both infarct volume and body weight loss. Furthermore, we show that serum NfL outperformed infarct volume in predicting a good outcome in stroked rats. These correlations were supported with evidence that NSC EV, a robust biologic therapy for AIS, significantly reduces serum NfL levels as well as preserves brain tissue and neurologic function. Overall, these data illustrate that serum NfL may be a useful biomarker to assess the therapeutic potential of investigational products for AIS to be directly translated to successful AIS clinical trials.

## Availability of data and materials

Data generated from the current study is available from the corresponding author upon reasonable request.

## Abbreviations

*AIS*: acute ischemic stroke
*ALS*: amyotrophic lateral sclerosis
ANOVA: analysis of variance
*ATP*: adenosine triphosphate
*AUC*: area under curve
*CCA*: common carotid artery
*CCL2*: C-C motif chemokine ligand 2
*ECA*: external carotid artery
*EV*: extracellular vesicle
*EVT*: endovascular thrombectomy
HLVe: hemispheric lesion volume with edema correction
*ICA*: internal carotid artery
*ICAM-1*: intercellular adhesion molecule 1
*IFNY*: interferon-gamma
*IL-1β*: interleukin-1 beta
*IL-6*: interleukin-6
*MCA*: middle cerebral artery
mRS: modified rankin scale
*NfL*: neurofilament light chain
NIH: National Institutes of Health
NIHSS: NIH stroke scale
*NSC*: neural stem cell
*ROC*: receiver operating characteristics
*S100B*: S100 calcium binding protein
SEM: standard error of the mean
*TBI*: traumatic brain injury
TFF: tangential flow filtration
*tMCAO*: transient middle cerebral artery occlusion
*TNFα*: tumor necrosis factor-alpha
*TTC*: triphenyl tetrazolium chloride

## Acknowledgements

We thank the University of Georgia University Research Animal Resources staff for their animal husbandry assistance.

